# NRL- and CRX-guided gene network modulates photoreceptor presynapse size and positioning during retinal development

**DOI:** 10.1101/753012

**Authors:** D. Thad Whitaker, Anupam K. Mondal, Hannah Fann, Passley Hargrove, Matthew J. Brooks, Vijender Chaitankar, Wenhan Yu, Zhijian Wu, Soo-Young Kim, Anand Swaroop

**Affiliations:** Neurobiology, Neurodegeneration & Repair Laboratory, National Eye Institute, National Institutes of Health, Bethesda, Maryland; Texas A&M Institute for Neuroscience, Texas A&M University, College Station, Texas; Institute of Biomedical Sciences, George Washington University, Washington, District of Columbia; Ocular Gene Therapy Core, National Eye Institute, National Institutes of Health, Bethesda, Maryland

**Keywords:** RNAi screen, neuronal differentiation, synaptogenesis, morphogenesis, regulatory network

## Abstract

Unique morphologies of rod and cone photoreceptor presynaptic terminals permit the formation of synapses onto interneurons during retina development. We integrated multiple “omics” datasets of developing rod and S-cone-like photoreceptors and identified 719 genes that are regulated by NRL and CRX, critical transcriptional regulators of rod differentiation, as candidates for controlling presynapse morphology. *In vivo* knockdown of 72 candidate genes in the developing retina uncovered 26 genes that alter size and/or positioning of rod spherules in the outer plexiform layer. Co-expression of seven cDNAs with their cognate shRNAs rescued the rod presynapse phenotype. Loss of function of four genes in germline or by an AAV-mediated CRISPR/Cas9 strategy validated RNAi screen findings. A protein interaction network analysis of the 26 positive effectors revealed additional candidates in the NRL/CRX-regulated presynapse morphology-associated gene network. Follow-up knockdowns of two novel candidates support the proposed network. Our studies demonstrate a requirement of multiple components in a modular network for rod presynapse morphogenesis and provide a functional genomic framework for deciphering genetic determinants of morphological specification during development.

**Author Summary:** The relationship between neuronal morphology and function has been recognized for over 100 years. However, we still have poor understanding of genes and proteins that control morphogenesis of a specific neuron. In the current study, we address this connection between gene expression and neural morphology by identifying and knocking down a subset of expressed genes in rod photoreceptors. We ascertained a number of candidate genes controlling photoreceptor pre-synaptic terminal morphology, which is necessary for its connection with second-order neurons in the retinal circuit. Furthermore, we have curated a more plausible network of genes, either identified in our study or predicted, that are enriched for processes underlying photoreceptor morphogenesis. We suggest that our work will provide a framework for dissecting genetic basis of neuronal architecture and assist in better design of cell replacement therapies for retinal degeneration.

## Introduction

The human central nervous system contains approximately 86 billion neurons that form 150 trillion synapses (1, 2). Each neuronal cell type possesses a stereotyped morphological standard, creating precise synaptic connections. Over 100 years ago, Santiago Ramón y Cajal predicted intimate structure-function relationships within neural circuits based on elaborate observations in the retina and the brain (3). The developmental trajectory of a neuron constricts its essential features towards that of its class to ensure a typical morphology and function, which are stringently and largely controlled by transcriptional control mechanisms (4–8). Genetic determinants that define neuronal size, shape, and structure are often unclear, and the mechanisms underlying the precise synaptic organization are poorly understood.

The retina offers an accessible and relatively less complex paradigm to delineate genetic underpinnings of intricate links between form and function. Development of the retina and associated visual circuits are remarkably conserved during evolution (9). The mammalian retina possesses five major classes of neurons that contribute to overall synaptic circuitry and organization (10): photoreceptors, bipolar cells, horizontal cells, amacrine cells, and ganglion cells (Fig. 1A), and each class consists of between two to 70 cell types (11). The two types of photoreceptors, cones and rods, initiate the visual process. Cones mediate high resolution day-light vision and permit color discrimination via distinct visual pigments; in contrast, rods possess a single visual pigment (rhodopsin) and carry out dim-light vision because of high sensitivity of photon capture (12, 13). Ultraviolet and short-wavelength (S-, “blue”) cones were the first photoreceptors to evolve in vertebrates yet rods dominate the retinal landscape in mammals (14, 15). Genetic and lineage-tracing studies suggest that most, if not all, rod photoreceptors in the mouse retina originate from S-cones (16). Both cone and rod photoreceptors synapse onto bipolar and horizontal cells in the outer plexiform layer (OPL) using a specialized ribbon organelle for sustained and graded transmission of visual signals (17, 18). Nonetheless, cones and rods exhibit distinct presynaptic morphologies, which are critical for their unique physiological characteristics (10, 13, 19). Cones possess a large presynaptic terminal (called “pedicle”) with multiple ribbon synapses, forming complex circuits in the OPL with multiple interneurons (20). In contrast, the rods have a relatively small presynaptic terminal (called “spherule”) with a single ribbon and connect to only a single class of bipolar neuron and the horizontal cell, illustrating a simpler neuronal circuit (21).

**Figure 1.**
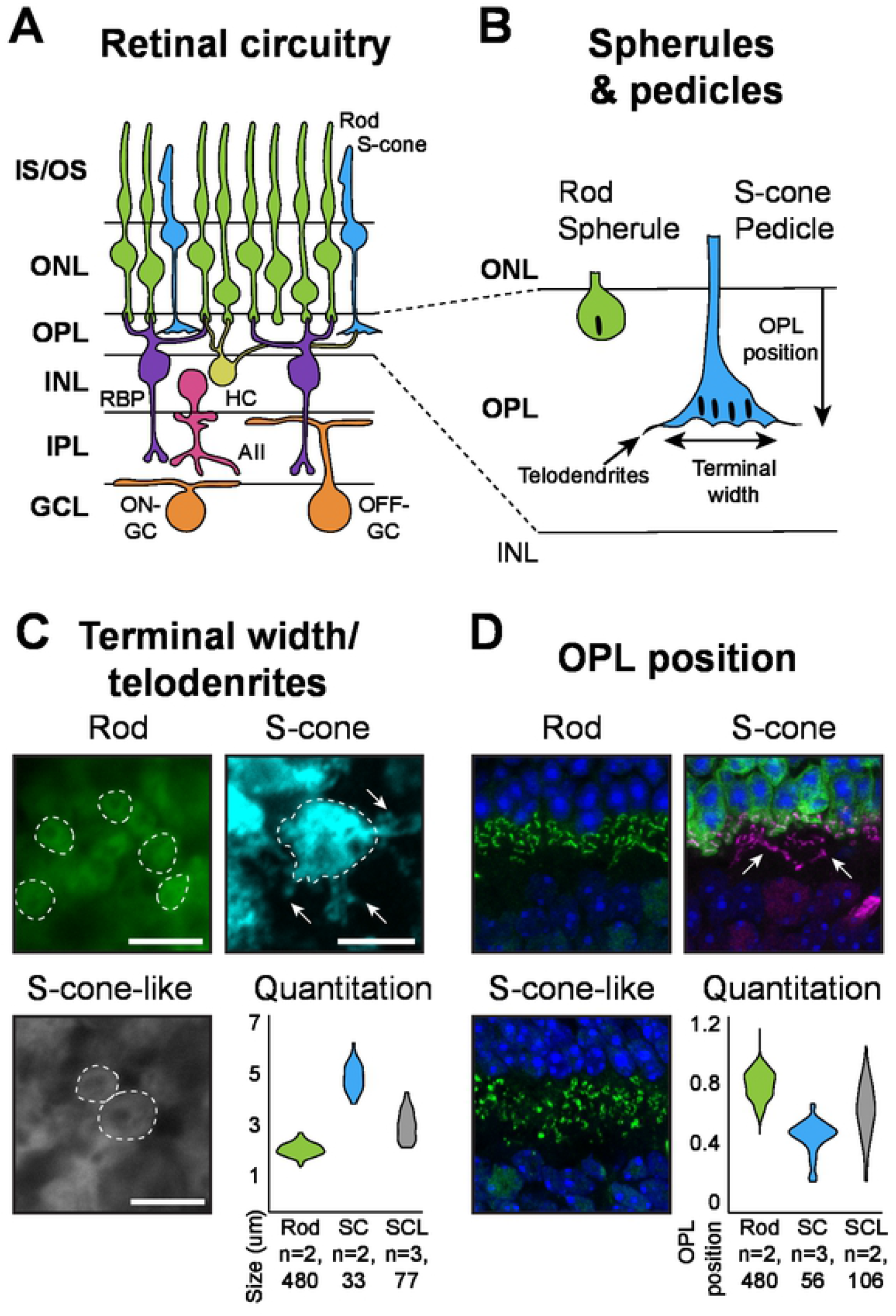
Assays for evaluation of morphology of rod and cone photoreceptors. (A) Schematic of neuron circuitry in the mammalian retina. Major cell types in the rod photoreceptor pathway are shown. IS/OS: inner/outer segments; ONL: outer nuclear layer; OPL: outer plexiform layer; INL: inner nuclear layer; IPL: inner plexiform layer; GCL: ganglion cell layer; RBP: rod bipolar cell; HC: horizontal cell; AII/A17: amacrine cells; GC: ganglion cell. (B) Cartoons showing features of a rod spherule and cone pedicle. The presynaptic terminals of rod and cone photoreceptors reveal three distinct morphological characteristics: presence/absence of telodendrites, size of terminals, and the sublaminal position within the OPL. (C) Spherules and pedicles are outlined. Width was calculated by defining the area and then calculating the diameter. Width of S-cone and S-cone-like pedicles are larger than spherules. S-cones also show noticeable telodendrites (arrows). (D) Sub-laminal position of presynaptic terminals in OPL. Terminals were defined as rod, cone, or cone-like using the *Nrlp*-GFP and *Nrlp*-GFP;*Nrl^−/−^* reporter mice. Ribbon position was used as a proxy for terminal positioning. The rods synapse closer to the ONL, whereas S-cones (in this panel, rods are labeled with GFP and examples of S-cone ribbon position are marked with arrow) and cone-like cells penetrate further into the OPL. Positions in OPL: 0=INL; 1=ONL.

Induction of the <u>N</u>eural <u>R</u>etina <u>L</u>eucine zipper (NRL) transcription factor is both necessary and sufficient to actively transition developing S-cones to the rod cell fate (22, 23). NRL acts synergistically with homeodomain transcription factor CRX and multiple transcription-regulatory proteins to control global pattern of gene expression in rods (24, 25). The transcriptome of developing mouse rods exhibits a dramatic shift between postnatal day (P)6 to P10 (16), which encompasses the initial period of synaptogenesis (26). Thus, a subset of these genes is predicted to determine the formation of rod spherules and subsequently synaptic connections. Loss of *Nrl* in mice transforms the post-mitotic photoreceptor precursors fated to be rods to S-cone-like (SCL) photoreceptors (22), which possess a cone pedicle-like presynaptic terminal morphology and cone-like function (27, 28). We therefore surmised that the rod spherule morphology, positioning, and function are guided largely by NRL-transcriptional targets, which are induced or repressed in rods at or after P6 during mouse retina development.

To identify genetic determinants of the stereotypical structure and positioning of a rod spherule, we took advantage of the NRL-mediated gene network underlying rod differentiation and selected 72 potential candidate genes for *in vivo* RNAi screening, as successfully demonstrated recently by another group (29). Of these, knockdown of 26 genes in the mouse retina revealed an altered spherule size and/or location in the sublamina within the OPL, indicating a transition from rod to S-cone-like pedicle structure. We validated the RNAi screen by cDNA rescue experiments and loss of function studies in mice. A NRL- and CRX-guided protein interaction network, developed from the 26 positive effectors, identified additional candidates associated with presynapse morphology and positioning; of these, two candidates that were tested further substantiated our findings. Our studies thus utilize “omics”-driven phenotype screening in mammalian retina to dissect the complex process of neuronal morphogenesis during development.

## Results

### Assay parameters to examine presynaptic structure and location

Using fluorescence reporters for *in vivo* electroporation in the mouse retina, we can identify at least three discrete gross morphological features at the photoreceptor presynaptic terminal differentiating between rod and cone presynapse terminals. These include: overall width, size, or diameter of the terminals; sublamination pattern in the OPL; and the presence of telodendrites (Figs. 1A, B, S1). The width of rod spherules is much smaller than either S-cone or SCL pedicles in the *Nrl^−/−^* retina (rods, 2.091±0.011 μm; S-cones, 4.917±0.092 μm; SCL, 3.036±0.063 μm, mean±SEM), and rods do not project telodendrites from the terminal body as in cones (Fig. 1C). In the OPL of wild type mouse retina, rod and cone terminals exhibit distinct lamination, with spherules synapsing close to the photoreceptor cell bodies and pedicles closer to the bipolar cell nuclei. To quantify the distinction, we calculated relative distance by assigning 0 to 1.0 position within the OPL, where 0 corresponds to synaptic terminal location at the top of bipolar nuclei and 1.0 to the bottom of photoreceptor nuclei (Fig. S1A). Using these criteria, the relative distance for rod spherules and S-cone pedicles was measured at 0.796±0.005 and 0.449±0.015 (mean±SEM), respectively (Fig. 1D). The terminals of SCLs in the *Nrl^−/−^* retina were more widely distributed throughout the OPL at relative distance of 0.628±0.127. The anatomical position of the rod photoreceptor did not influence any of the presynapse structural features (Fig. S1). These data are consistent with other reports (30, 31) and provide a baseline for assaying divergent features between rods and cones after genetic or experimental manipulations.

### Candidate determinants of rod spherule morphogenesis

We initially focused on identification of genes that modulate two presynaptic features: width/size and the relative position in the OPL. Taking advantage of the published transcriptomes of developing rods and *Nrl^−/−^* SCLs (16) and of the ChIP-seq datasets for NRL and CRX (32, 33), we identified 719 potential candidate genes by using the following filtering criteria (Fig. 2): (1) a minimum threshold of 25 counts per million (CPM) to reflect high expression in rods at any developmental stage; (2) increase in expression after P6, concurrent with rod presynapse morphogenesis and positioning; (3) at least two-fold enrichment in rods compared to SCL photoreceptors; and (4) direct transcriptional targets of NRL and CRX, as suggested by the ChIP-seq data (Fig. S3). For identifying genes associated with presynapse morphology, we selected 72 candidates (Table S1) using a combination of gene ontology (GO) analysis and literature search for understudied (no or few citations) genes.

**Figure 2.**
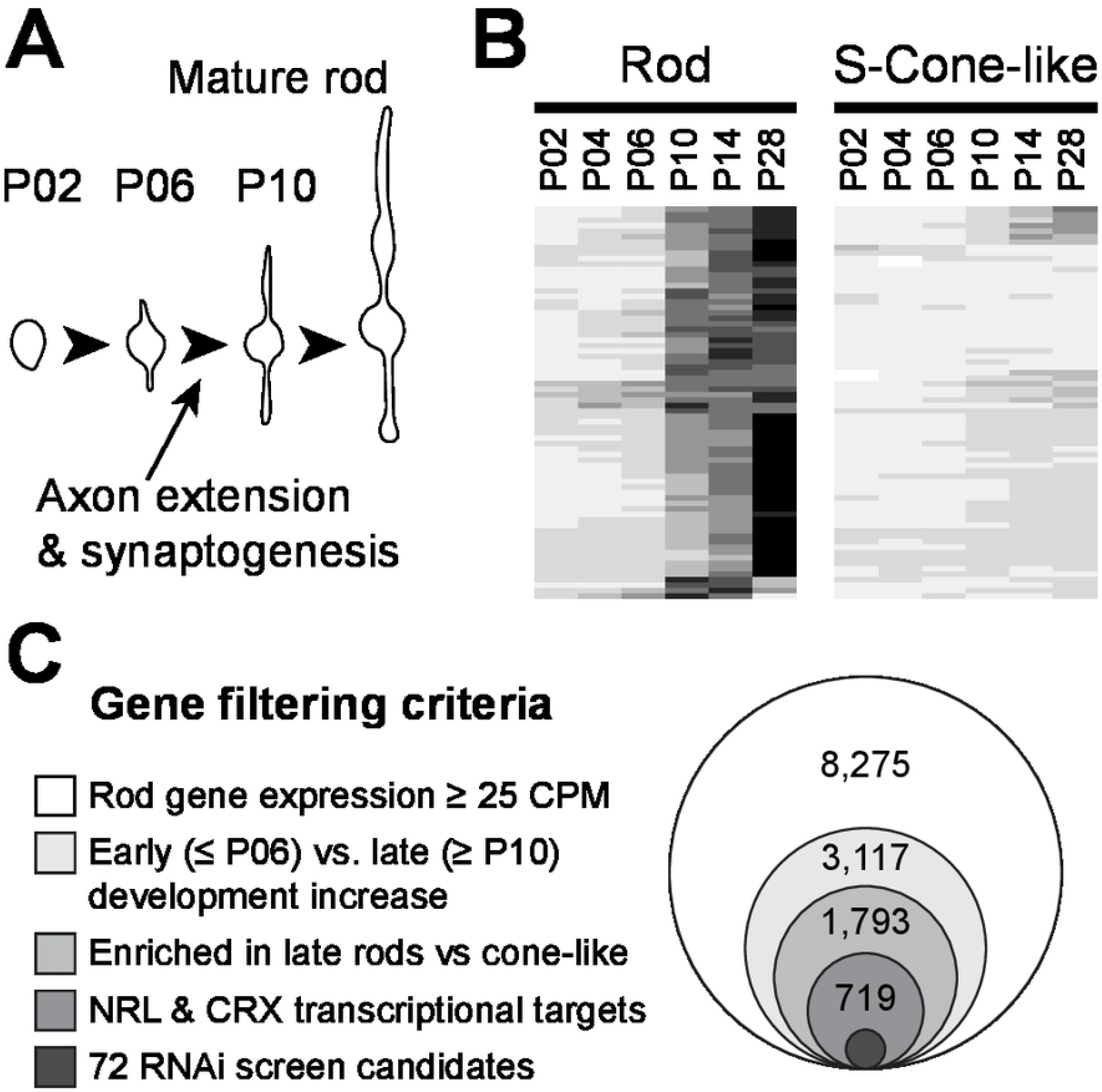
Identification of candidate RNAi screen genes. (A) Schematic of rod morphology throughout development. Key neurite and synapse development occurs during P6 – P10 (arrow), the same time the rod transcriptome is undergoing major shifts. (B) Developmental heat map of gene expression between rod and S-cone-like cells. Z-score normalized gene expression of all 72 genes selected shows dramatic shift in expression between P6 and P10 in rods but not cone-like cells. (C) Gene filtering paradigm. Using available “omics” data, we narrowed total transcriptome down to 72 candidates for the current screen.

### RNAi screen and phenotype rescue for presynaptic terminal width/size

Given that the “default” S-cone photoreceptors (25) can be converted to rods by NRL (23), we reasoned that the genes associated with rod morphogenesis restrict the pedicle size to that of the spherule and their loss of function would generally increase rod spherule width. We therefore performed electroporation of short hairpin RNAs (shRNAs) against 72 candidate genes in addition to relevant controls into P0-P1 mouse retina *in vivo*. We also included fluorescent reporter plasmids to sparsely label electroporated rod photoreceptors. The retinal sections were assayed for spherule width 3-weeks after electroporation. In screened control mice, our measurements of terminal size (1.8 to 2 μm; Fig. S1) corresponded well with the spherule width of uninjected retinas (Fig. S2). Quantitative evaluation of seven out of the 72 assayed genes (*Dpf3, Epb4.1l2, Grtp1, Kcnj14, Llgl2, Rab28*, and *Rom1*) significantly increased the width of rod spherules upon RNAi knockdown (Fig. 3). Interestingly, knockdown of six other genes (*Alpl1, Asrgl1, Bbs9, Eif3f, Plcd3, Plch2*) revealed smaller spherules (Table S1), suggesting an intrinsic and stringent synergistic and antagonistic control of presynapse terminal morphology to refine the size of the rod spherule.

**Figure 3.**
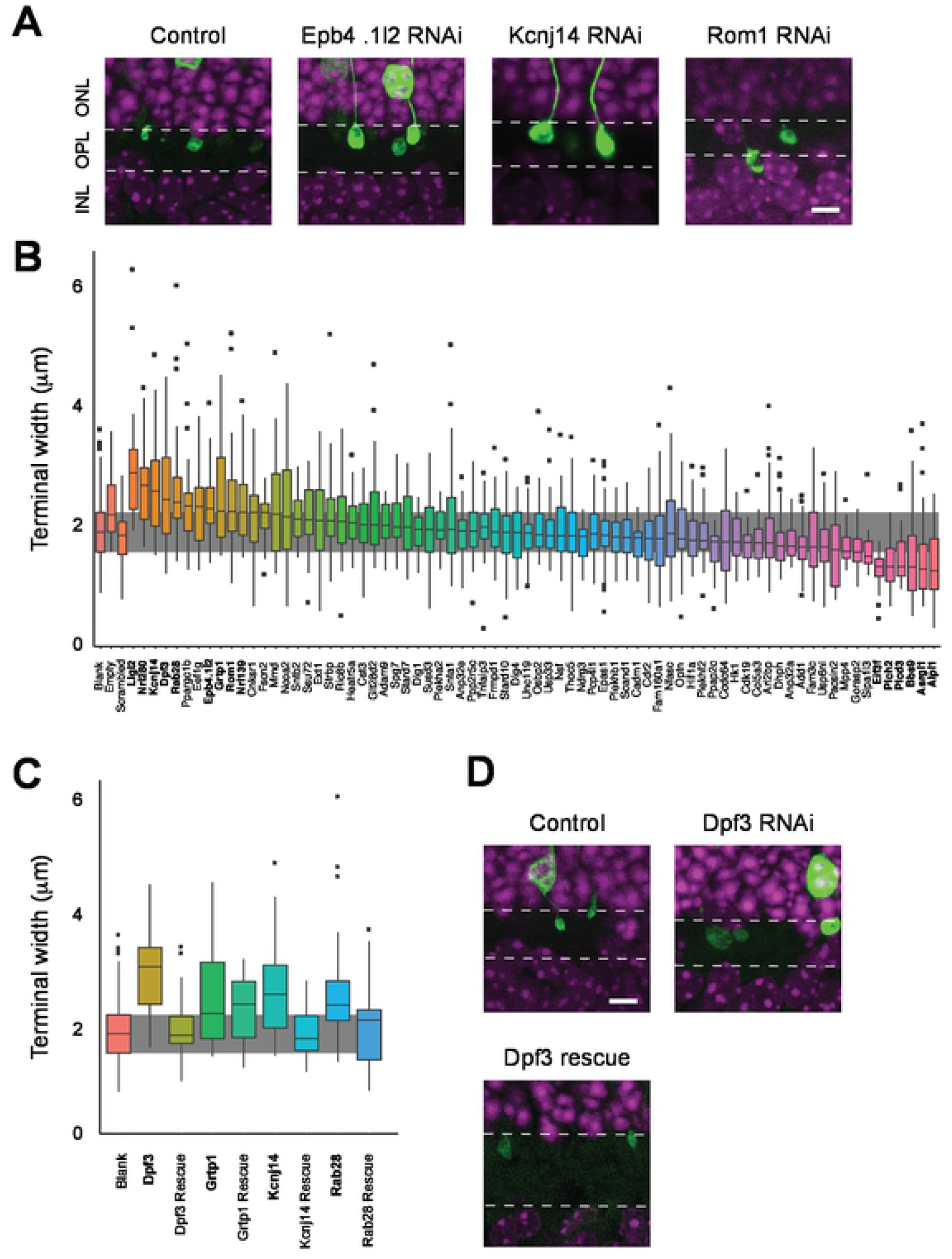
RNAi screen for presynaptic terminal size. (A) Representative images of three RNAi knockdowns showing differential terminal size compared to control. Scale bar = 5μm. (B) Quantitative analysis of the RNAi screen. The graph is organized with the three independent controls (reporter only, scrambeled shRNA, and empty vector, respectively) on the left, and then 72 gene-specific knockdowns arranged in order of increasing median of terminal size. Grey background represents the interquartile range (IQR) of the reporter-only control. The genes in bold show significantly different size compared to the controls. (C) Rescue of the RNAi phenotype by reintroduction of modified cDNAs designed to circumvent RNAi inhibition (see Fig. S4). All four rescue RNAi experiments evaluated demonstrated a recovery of the original spherule size. Control data reproduced from Figure 3B. The genes in bold show significantly different size compared to the controls. (D) Representative images for one rescue experiment. *Dpf3* knockdown results in enlarged terminals; re-addition shows a complete recovery of the rod spherule phenotype. Scale bar = 5μm. n≥25 spherules per construct were quantified for each experiment.

To validate RNAi screen and reject type I errors, we performed rescue of the spherule phenotype by coexpressing the cDNA of the targeted gene with modifications in the shRNA-binding site (Fig. S4) with the original shRNA. Introduction of four cDNAs (*Dpf3, Grtp1, Kcnj14*, and *Rab28*) together with their respective RNAi molecule prevented or reduced the increase in spherule width that was observed with RNAi alone, demonstrating the target specificity of the evaluated knockdowns (Fig. 3C,D; Table S1).

### RNAi screen and phenotype rescue for spherule positioning in OPL

We then examined the OPL sublamina location of the rod spherules using the same strategy. The relative distance of the spherule from the top of bipolar cell nuclei provided measurements similar to the ribbon location as determined by Ribeye staining (Figure S2). ShRNA-knockdown of 23 out of 72 genes tested by electroporation in the mouse retina impacted the spherule location in the OPL (Fig. 4A,B,C; Table S1). Of these, 14 gene knockdowns resulted in patterns similar to native S-cones (e.g., *Kcnj14*), and nine additional genes deviated significantly from rod spherule controls with an intermediate sublamina location between the rod and cone terminals (e.g., *Ncoa2*). Both spherule width and sublamina location were affected by knockdown of four genes – *Epb4.1l2, Kcnj14, Llgl2*, and *Rom1* (Figs. 3B, 4C).

**Figure 4.**
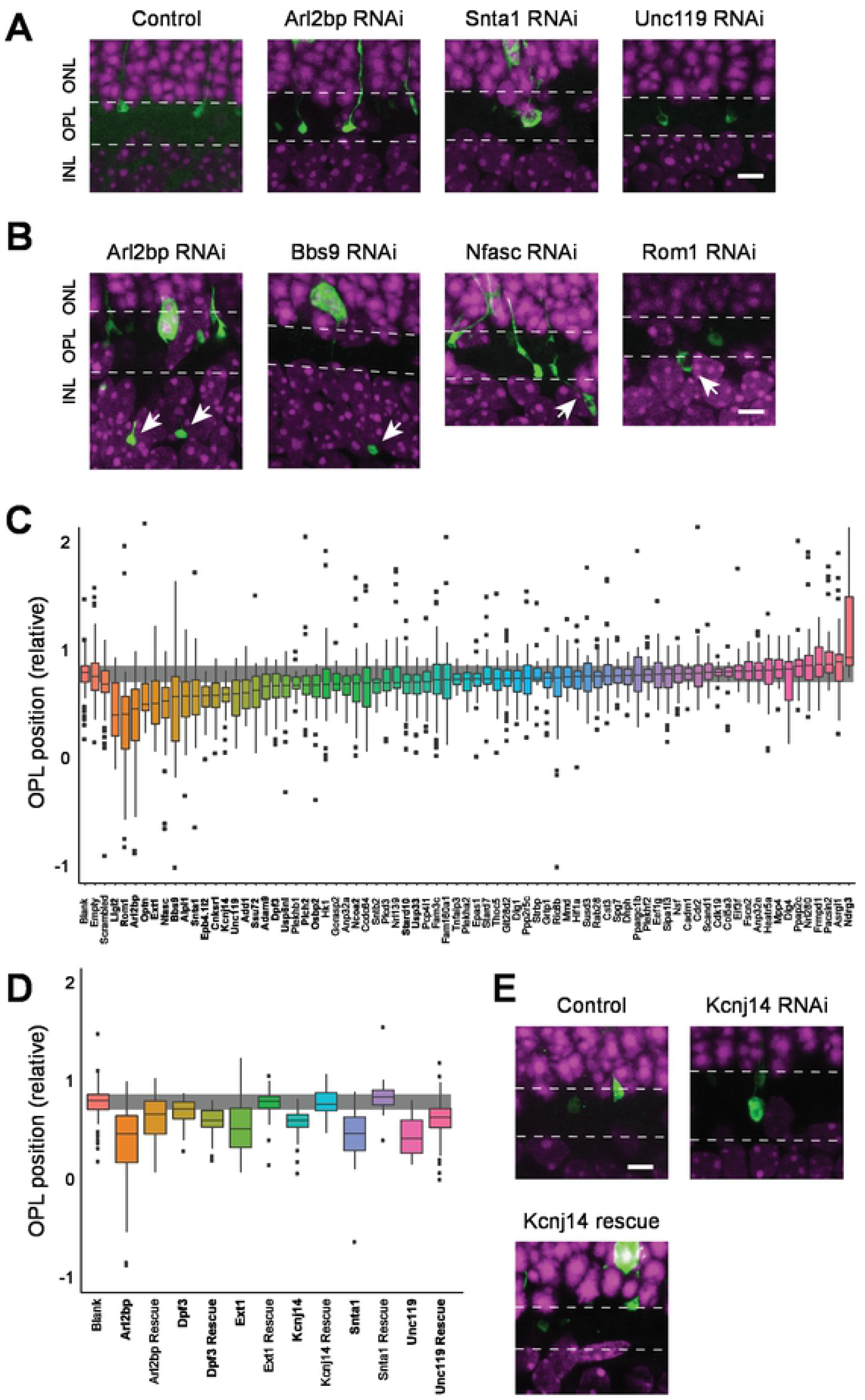
RNAi screen for determinants of sub-lamina position. (A) Representative images of three RNAi knockdowns, showing differential position of terminals in the OPL. Scale bar = 5μm. (B) Examples of severe terminal positioning defects observed after RNAi knockdown in some instances (represented with negative position values in Fig. 1C). The axon/terminal extended beyond the OPL and into the INL in these examples. Scale bar = 5μm. (C) Quantitative analysis of the RNAi screen. Graph is organized with the three independent controls (reporter only, scrambeled shRNA, and empty vector, respectively) on the left, and then 72 gene-specific knockdowns arranged in order of decreasing median of sublamina position in OPL. Grey background represents the interquartile range (IQR) of the reporter-only control. The genes in bold show significantly different positioning compared to the controls. (D) Rescue of the RNAi phenotype by addition of modified cDNAs. Four of six rescue experiments demonstrated a recovery of the original spherule position. *Unc119* showed a partial recovery. Control data reproduced from Fig. 4C. The genes in bold show significantly different positioning compared to the controls. (E) Representative images for *Kcnj14* rescue experiment. *Kcnj14* knockdown resulted in altered positioning of the presynaptic terminal; re-addition of the modified cDNA showed a complete recovery of the rod spherule phenotype. Scale bar = 5μm. n≥25 spherules per construct were quantified for each experiment.

Altered OPL positioning phenotypes detected by RNAi could be rescued by concomitant injection of cDNAs for four genes – *Arl2bp, Ext1, Kcnj14*, and *Snta1* (Fig. 4D,E; Table S1). Curiously, *Dpf3* cDNA prevented the shRNA-mediated increase in spherule size but exacerbated the OPL position phenotype. The introduction of *Unc119* cDNA lessened the impact of RNAi on the OPL position. These data validated the knockdown specificity for synapse location and our overall approach for identifying genes associated with spherule morphogenesis.

### Altered spherule size by germ-line or AAV-mediated loss of function in mouse retina

We then obtained previously-established mouse strains with germ-line loss of *Epb4.1l2* (34) or *Snta1* (35). In addition, we generated loss-of-function (LOF) of *Dpf3* and *Llgl2* in mouse retina by an AAV8-mediated CRISPR/Cas9 strategy, as described previously (36). Morphology assays of rod spherules in the retina of all four mouse models demonstrated a significant trend towards larger terminal size (Fig. 5). Unlike a previous report (34), we did not see retracted spherules in the *Epb4.1l2* LOF rod photoreceptors, possibly due to the differences in ages between the two studies. Spherule size was significantly enhanced in *Snta1^−/−^* retina even though the RNAi screen revealed a much smaller difference in size but a more pronounced change in terminal location (see Figs. 4, 5). Additional *Snta1*-knockdown screens using RNAi electroporation in the mouse retina validated the results from *Snta1^−/−^* mice and demonstrated a subtle increase in terminal size (data not shown), consistent with the knockout retina. The presynapse location of rod spherules did not seem to be altered in the retina of four mouse models (Fig. 5).

**Figure 5.**
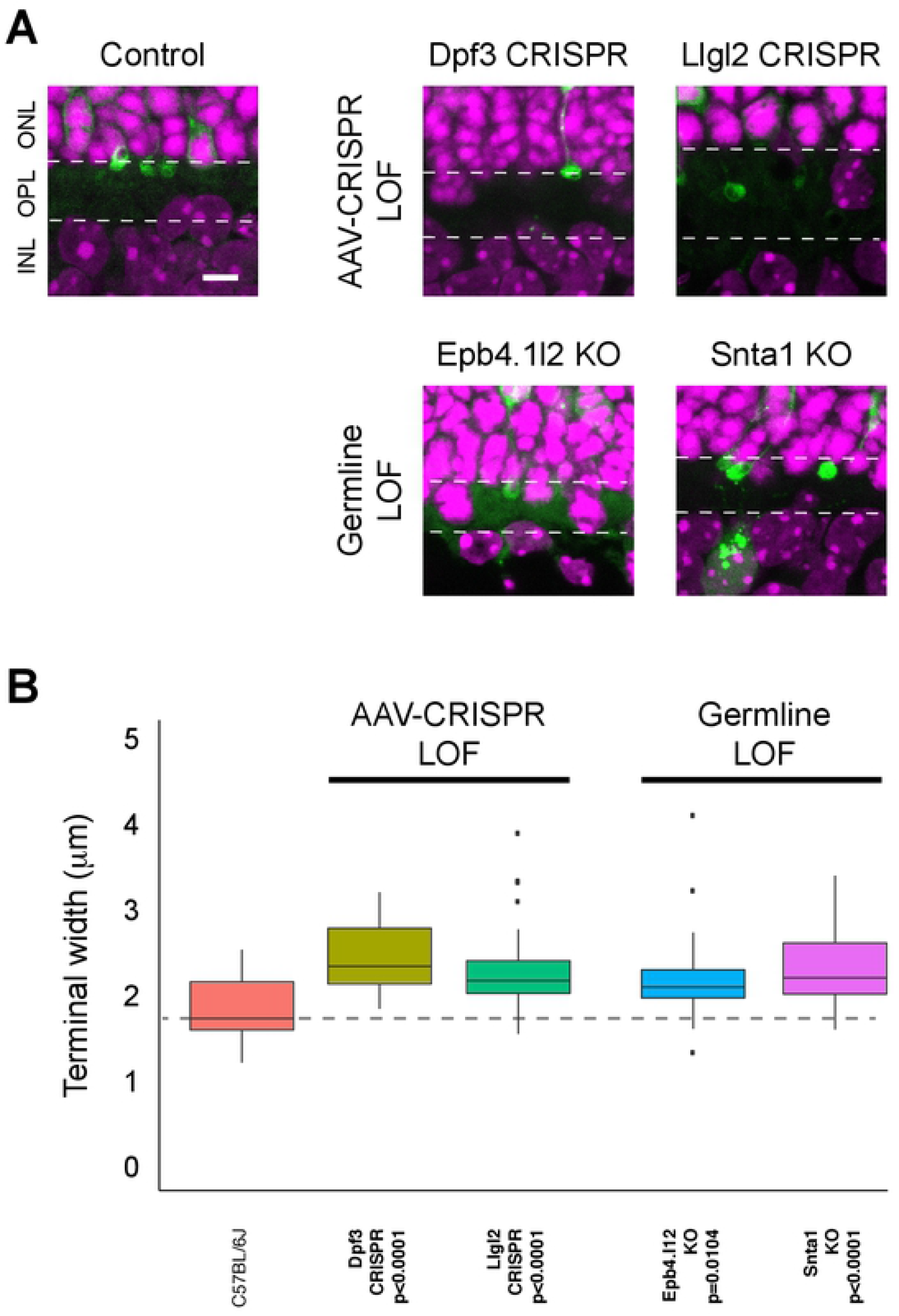
Loss of function rod photoreceptors recapitulate enlarged terminals. (A) Representative images of the enlarged spherule terminals in four LOF models, generated either by CRISPR/Cas9 delivered with AAV8 or by fluorescent labeling of the genetic null (knockout, KO) mouse strains. All four LOF models displayed enlarged spherule sizes compared to controls. Scale bar = 5μm. (B) Quantitative analysis of spherules for terminal width in wild type and LOF models. All four models tested had somewhat enlarged spherules. Dashed line indicates median of control. Scale bar: 5μm. n≥25 spherules per construct were quantified for each experiment.

### Deciphering presynapse protein network and follow-up validation studies

To advance our findings and identify additional components associated with spherule morphology, we constructed a protein interaction network using STRING (37) from our curated list of 719 genes (Fig. 2) and their high-confidence first-order neighbors (Fig. S5). We mapped the shortest path connections for all pairwise combinations of the 26 proteins corresponding to genes that demonstrated directionality away from the rod spherule and towards the cone-like pedicle phenotype upon knockdown, revealing 8,443 possible pathways (Fig. 6). We further prioritized the paths to contain at least one other experimentally validated protein, while not including any “negative” node, i.e., genes that exhibited no pedicle-like features after knockdown. The qualifying 96 shortest paths form a network including 71 total proteins that span 17 experimentally-identified “positive effector” nodes; these paths are enriched for synapse-associated ontology terms (Figs. 6C, S5D). We predict that several proteins in these interaction pathways would contribute to spherule formation, and together with their cognate interactors, represent the gene network underlying photoreceptor presynapse morphogensis (hereafter termed <u>P</u>resynapse <u>M</u>orphology <u>A</u>ssociated Network, PMAN; Figure 6B). Two observations were apparent from PMAN; (i) the capture of primary photoreceptor transcription factor interactions (*Nrl-Crx-Nr2e3*) in a single anticipated pathway, and (ii) a strategic placement of *Ncoa2* based on its high degree of connectedness and betweeness-centralitiy, implicating its importance in the network (Fig. 6B).

**Figure 6.**
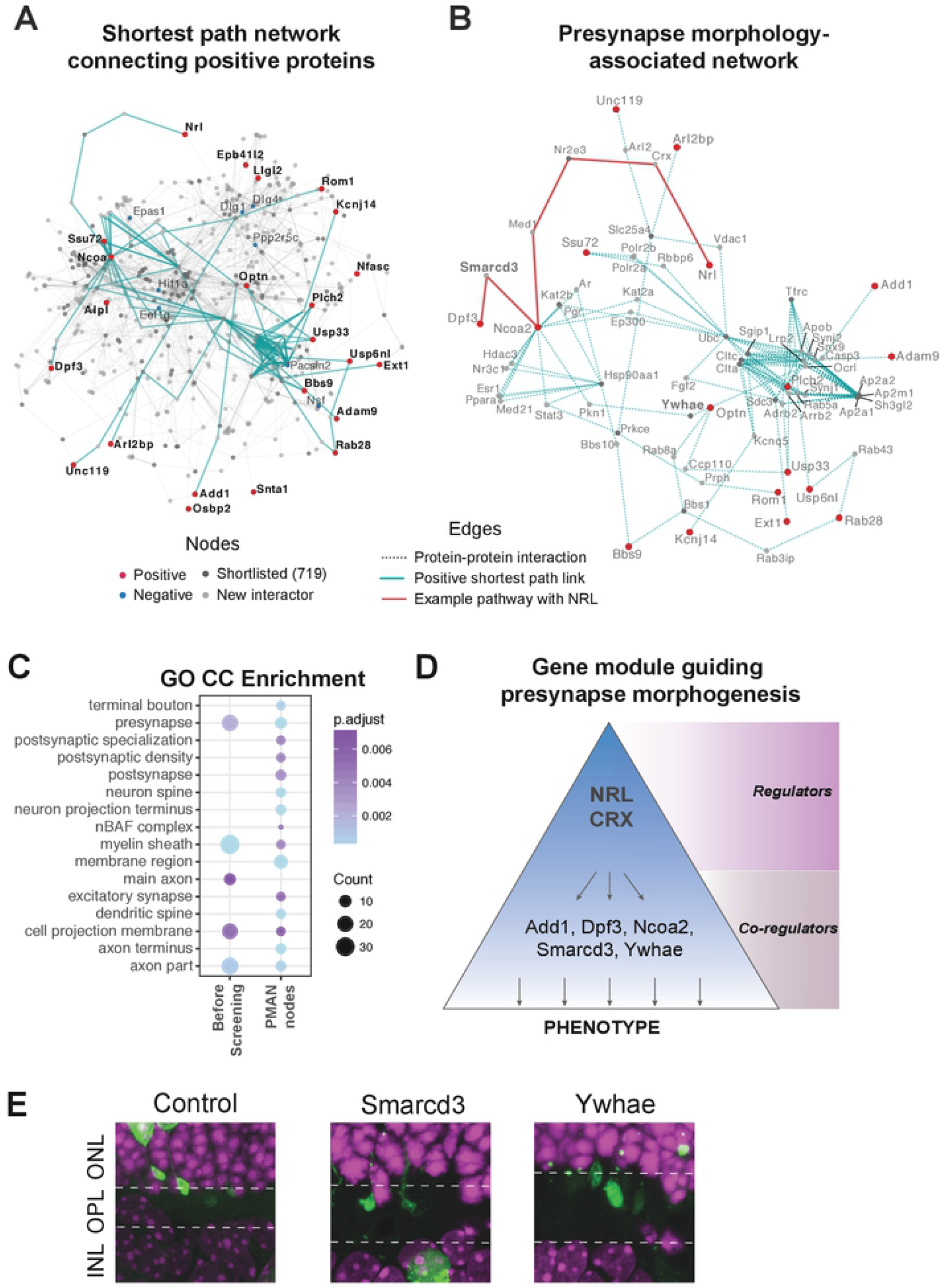
Construction of <u>p</u>resynapse <u>m</u>orphology <u>a</u>ssociated gene <u>n</u>etwork (PMAN) and experimental validation of two candidates. (A) A network of shortest paths connecting every pair of experimentally-verified gene nodes. Genes with any synaptic phenotype (“Positive”) identified in the RNAi screen are labeled in red in the network. Targets showing no phenotype (“Negative”) are labeled in blue. Dark grey were genes in the initial candidate list, while others are colored grey. The lines connecting nodes represent protein-protein interactions as described in STRING. Teal edges highlight links in prioritized shortest paths with positive intermediates. (B) High confidence shortest path interactions in PMAN of filtered links connecting pairwise combinations of validated genes. Highlighted red is the shortest path connecting NRL to this network. Two new nodes, highlighted in bold (Smarcd3 and Ywhae), were selected for further validation. (C) Comparative GO (cellular component) enrichment in PMAN nodes and initial list of genes focusing on synapse specific GO terms. (D) A proposed model of tiered molecular network driving spherule morphogenesis. (E) RNAi knockdown of *Smarcd3* and *Ywhae* (selected from PMAN), showing aberrant spherule morphology. Scale bar: 5μm

To experimentally evaluate the network, we selected two candidate genes, *Smarcd3* and *Ywhae*, from the PMAN shortest paths (Fig. 6B). *In vivo* RNAi knockdown of these two genes in the developing mouse retina demonstrated aberrant rod spherule morphology (Fig. 6E). The significance of PMAN is highlighted by the capture of *Smarcd3*, a novel candidate that was not present in the initial shortlist of 719 candidates. These studies provide strong support to the premise of our protein interaction network and allow us to propose a modular organization of the gene regulatory network controlling presynapse development. With regulators such as NRL and CRX at the top of the hierarchy as primary drivers of rod differentiation, the presynapse terminal phenotype is guided through key co-regulators (e.g. *Add1, Dpf3, Ncoa2, Ywhae*) to produce precise native spherule structures (Fig. 6D).

## Discussion

Complex information processing in the nervous system is made possible by distinctive cellular architecture and highly-organized circuitry of neurons. How diverse morphologies of specialized neurons are acquired is poorly elucidated at the molecular level. Development of synaptic circuitry is hardwired into the genetic program (38). Therefore, a comprehensive delineation of connectivity between one population of neurons to another would require a better understanding of gene regulatory networks that control appropriate localization and structure of axon boutons and dendritic branches during synaptogenesis and for neurotransmission. In the OPL of the retina, rod and cone photoreceptors construct the first visual synapse with bipolar and horizontal neurons. Recent studies have begun to unravel the structural and molecular underpinnings of the photoreceptor ribbon synapse (39, 40). In addition to Ribeye (41) and Piccolino (a splice variant of Piccolo) (42), a few molecular components have been associated with precise photoreceptor presynapse morphology and wiring with bipolar cells; these include scaffold protein 4.1G (also called Epb4.1l2), cell adhesion protein ELFN1 (43), and extracellular proteins Pikachurin (44) and auxillary calcium channel subunit α2δ4 (45). Here, we define a set of 26 new molecular determinants that limit the size and/or axonal positioning of the spherule in the appropriate sublamina, two essential features associated with functional specificity of the rod photoreceptor synapse. We predict that augmented expression of these genes in cones would produce rod-like terminals. Almost one-third of the candidate genes selected after gene filtering demonstrated an altered spherule phenotype by RNAi-knockdown *in vivo.* We validated our findings by both rescue experiments using modified cDNAs and *in vivo* LOF models in mice. Furthermore, we suggest a modular gene network PMAN that controls rod spherule structure in developing retina.

*In vivo* electroporation has become a widely used technique in retinal biology for evaluation of photoreceptor phenotypes (29, 46, 47). The variables including age, litter size, sex and precise site of injection have not been shown to affect the phenotypes we studied in this report. As only actively dividing cells are primarily transfected by electroporation, we are evaluating a subpopulation of rod photoreceptors that are still early in their developmental trajectory. Transfection of one rod photoreceptor with reporter and RNAi molecule is independent of all neighboring cells. Our analysis therefore captures a large number of independent events in one experiment.

Complex morphogenic processes, such as acquisition of unique presynaptic structure, are predicted to involve multiple coordinately-regulated pathways and numerous specific components. We focused on NRL-mediated transcriptome changes because of its key role in controlling rod cell fate and function and selected only genes that were also regulated by CRX because of the synergy between NRL and CRX in activating rod differentiation (48). Furthermore, CRX has recently been shown to regulate genes associated with active zone formation in the photoreceptor presynapse (49). In addition to validating this fundamental concept, our knockdown screen implicates combinatorial contributions (synergistic and/or antagonistic) of a diverse array of genes in a modular fashion to ensure normal spherule architecture. For example, seven gene knockdowns resulted in enlarged rod spherules, with only two *Llgl2* and *Kcnj14* yielding terminals closer in size to S-cone-like pedicles of the *Nrl^−/−^* retina. Of the six gene knockdowns that resulted in significantly smaller spherules, three (*Alpl1, Bbs9*, and *Plch2*) demonstrated deeper penetration of spherules into the pedicle sublamina of the OPL. Interestingly, a total of 23 genes appear to control sublamina position of photoreceptor synapse. We note that OPL patterning is also influenced by signaling from the post-synaptic rod bipolar neurons (29). Elucidation of intrinsic molecular components, i.e., the members of the PMAN, and their extrinsic regulators will permit us to investigate how photoreceptors integrate in the neural circuitry of the developing retina (50), which in turn might help in designing better photoreceptor replacement strategies for retinal degenerative diseases. Given the key roles of NRL and CRX in rod photoreceptor development, we speculate that phenotypes beyond the two assayed here are regulated by the genes studied here. Indeed, knockdown of some of the genes also resulted in additional phenotypes (see Table S1); however, these morphological changes were not investigated in detail.

Our approach to integrate knockdown screens with protein interactions, followed by network analyses and prioritization, has allowed us to trace the connections and directionality of morphogenesis-associated genes. The resulting network recapitulates previously-reported interactions of photoreceptor genes and identifies key moderators like *Ncoa2* that hold strategic positions between other experimentally-identified genes and NRL. We hypothesize a model of functional segregation in rod synaptogenesis where cell-fate determining factors (NRL and CRX) control morphology of rod photoreceptors including synaptic architecture with co-regulators such as *Ncoa2, Add1, Ywhae and Dpf3*.

Beginning with two “omics” datasets and empirically-designed phenotypic screening assays, we have been able to deduce the PMAN, a highly enriched network of synapse components, from the large developing transcriptome datasets. Through iterative phenotype screenings and genetic analysis, PMAN could yield a comprehensive molecular organization underlying photoreceptor presynapse morphogenesis. Another approach to integrate genetics with morphological differences and function is the use of recombinant inbred mouse strains by identifying quantitative trait loci driving structural distinctions. The mouse strains would also permit evaluation of functional implications of unique neuronal morphologies. Our studies thus illustrate a framework of *in vivo* screening combined with a systems-based network approach in deconstructing neural organization and phenotypes.

## Materials and Methods

### Mice

All procedures involving the use of mice were approved by the Animal Care and Use Committee of the National Eye Institute (ASP# 650). Nrlp-GFP and Nrlp-GFP;*Nrl^−/−^* mice were described previously (51). *Epb4.1l2* (34) and *Snta1* (35) loss of function strains were obtained from RIKEN (Tsukuba, Ibaraki, Japan) and Jackson Laboratories (Bar Harbor, Maine, USA), respectively. C57BL/6J and CD-1 IGS mice were purchased from Jackson Laboratories and Charles River Laboratories (Wilmington, Massachusetts, USA), respectively. All mice were euthanized at P21 – P22.

### *In vivo* electroporation

Transfection of retinal cells was performed as originally described (46). Briefly, fluorescent and the RNAi Consortium’s shRNA (see Table S1) plasmids (500 ng/ul and 1,200 ng/ul, respectively) were mixed with Fast Green dye (0.1% final concentration). Postnatal day (P)0 – 1 mice were anesthetized on ice until unresponsive to foot pad pinch. Eyelids were cleaned with 70% ethanol; fused junction of eyelid opened with 25-gauge needle. Small, shallow puncture was made in the sclera with 30.5-gauge needle. Approximately 0.2 – 0.3 microliters of plasmid solution injected into the subretinal space with a blunt-ended Hamilton syringe (Reno, Nevada, USA). Following injection, five electrical pulses (80V, 50ms, 950ms intervals) applied across the head with the positive electrode on the injected eye. Pups were placed onto a heated pad until warm, marked (methods below), then returned to the mother. Ketoprofen (2-5mg/kg body weight) was injected immediately following electroporation before placing back on the heated pad.

### Animal marking

To reduce animal usage, two methods of identification were used (tail clipping and tattooing) to identify which constructs were injected. The tip of the pup’s tail was removed with a new, sharp razor blade. For tattooing, a small amount of green tattooing paste (Fine Science Tools) was applied to the footpad(s) of the mouse, a 30.5-gauge needle was used to puncture the skin and insert a small amount of paste, then the footpad was cleaned. Tattoos could be discerned without loss of color for up to the 3 weeks of our study.

### Immunohistochemistry

Eyes used to stain for photoreceptor characterization were fixed for 1 hour in 4% paraformaldehyde (PFA) in 1x PBS. The anterior portions of the eye (cornea and lens) were removed after 10 – 20 minutes of this fixation period and eye cups returned to fixative. For sections, eyes were transferred into 10% and 30% sucrose (overnight each) before embedding in O.C.T. compound. Sections were cut at 12 - 14μm and mounted on SuperFrost+ charged slides. Slices were permeabilized and blocked in 5% normal donkey serum and 1x PBS with 0.3% Triton X-100 (PBST) for 30 – 60 minutes then incubated in primary antibody (1:1000) overnight at 4°C. Slides were washed 3 – 4 times in 1x PBST for 10 minutes each before incubating with secondary antibodies (1:1000) and DAPI for 2 hours at room temperature in the dark. Slides were washed again 3 – 4 times in 1x PBST, excess solution removed, and coverslipped using FluoroMount-G^™^ (ThermoFisher, Waltham, MA, USA). For whole mount retinas, tissue was fixed as above but the retina was also isolated from the eye cup before further fixative and staining. For whole retina staining, IHC proceeded as for sections except with longer incubation times (3-hour blocking; 3 days primary/secondary antibody (1:1000) at room temperature; 1-hour washes; final overnight wash before mounting). For the shRNA screen retinas, eyes were fixed for 3+ hours in 4% PFA; retinas were isolated again after 10 – 20 minutes before being returned to fixative. Isolated retinas were mounted in 7% low-melting point agarose and sectioned at 100μm using a vibratome. We stained these sections with DAPI at 1:2500 overnight, shaking. Sections were mounted and coverslipped as above.

### Confocal imaging

For all images in this study, we utilized one of two confocal microscope systems: Zeiss (Oberkochen, Baden-Württemberg, Germany) confocal LSM 700 and LSM 780 systems. For terminal width and telodendrite presence in wild type and *Nrl^−/−^* C57BL/6J mice, whole-mount retinas were used. To identify pure S-cone pedicles, we identified S-opsin positive cone outer segments that double labeled with cone arrestin in the far ventral retina where true S-cones are present. Cone arrestin filled the entire cell, including the pedicle, and these S-cones were followed from outer segment to the pedicles. Z-stacks were collected to ensure entire terminal was imaged; maximum intensity projections were created for quantitation. For the OPL position of wild type and *Nrl^−/−^* retinas, single plane images were taken of Ribeye, DAPI, and the *Nrlp*-GFP transgenic fluorescence. For the *Nrl^−/−^* retina, we utilized the Nrlp-GFP transgenic marker to identify cone-like pedicles and S-cones. Images for shRNA screen and validation quantitation were collected with constant objective magnification (63x) and image resolution (1236 × 1236) for strict consistency between images. Under this higher magnification, single (or neighboring) spherules were imaged using Z-stacks (step size 0.5μm) through the entire spherule to collect the widest and deepest penetrating points. A maximum intensity projection image of all stacks was created for downstream quantiation.

### AAV-mediated CRISPR knockout

The dual vector system of AAV-CRISPR was previously described (36) and maintained in the lab. Expression of SpCas9 was driven by a photoreceptor-specific rhodopsin kinase (RK) promoter (52). The sgRNA against *Dpf3* or *Llgl2* were selected using the online program developed by Dr. Feng Zhang’s laboratory (http://crispr.mit.edu/) and the Benchling (https://benchling.com/). The sequence encoding each sgRNA was cloned into an existing AAV shuttle vector maintained in the lab which contains the sgRNA scaffold with the U6 promoter. A tdTomato reporter gene driven by the RK promoter was also included in this vector to indicate AAV transduction.

SgRNA_Dpf3target: 5’-ACTGCCGGAGCTACAACTCG-3’ (PAM: 5’-AGG-3’)

SgRNA_Llgl2target: 5’-CAGAGCGGGTTCCGATGGCC-3’ (PAM: 5’-AGG-3’)

The AAV production was conducted as described (53). Briefly, HEK-293 cells (ATCC^®^ CRL-1573) were transfected with the ITR-containing vector plasmid carrying the spCas9 cassette or the sgRNA cassette, together with the two helper plasmids containing genes for viral replication/capsid assembly and adenoviral genes essential for AAV replication respectively.

The cells were collected after forty-eight hours and were disrupted by a microfluidizer (HC 2000; Microfluidics Corporation, Newton, MA). The cell debris were removed by centrifugation. Residual DNA was removed using Benzonase (100 U/mL). Vector particles were then concentrated using 8% polyethylene glycol (PEG) 8000 and further purified by Cesium Chloride (CsCl) density gradient centrifugation. The band containing the vector was collected with an 18-gauge needle, dialyzed against Tris-buffered saline (10mM Tris-Cl, 180mM NaCl, pH7.4) with 0. 001% Pluronic F-68, and stored frozen at −80°C. The vectors were titered by real time PCR using linearized plasmid standards and primers against the RK promoter.

To knock out *Dpf3* or *Llgl2* in mouse photoreceptors, the SpCas9 vector and the sgRNA vector were co-injected into subretinal space of C57BL/6J mice at P0-P1 as done in our *in vivo* electroporation experiments without subsequent electrical pulses applied.

### Synaptic terminal quantification

The ImageJ open-source software (54) was used for all synaptic terminal quantification. Images were randomly assigned an arbitrary number and terminals marked for future measurements. Blinded observers each made a series of three measurements for each terminal (shown in Figure S1A) using ImageJ’s line segment measurement tool. These were compared for consistencies between observers, and only minor, non-significant deviations seen. Of the 36 observations made in more than one retina, we found 27 (75%) showed no significant differences between retinas. A master table (Table S1) reports all averaged observations.

### Site-directed mutagenesis

Mouse full-length cDNAs were ordered from one of two companies (Dharmacon: Lafayette, Colorado, USA; Source BioScience: Nottingham, United Kingdom) in the pCMV-Sport6 expression construct. One clone (*Dpf3*) was originally in another construct and so was cloned into pCMV-Sport6 before further processing. We used the Q5^®^ Site-Directed Mutagenesis Kit (New England Biolabs: Ipswich, Massachusetts, USA) to create 5 – 7 synonymous mutations at the shRNA-binding site in the coding region or 5 – 7 nucleotide alterations for shRNAs that bound to the 3’ untranslated region (see Supplemental Figure 7 for exact nucleotide alterations). After ligation, plasmids were transformed into DH5*α*^™^ Competent Cells (ThermoFisher). Individual colonies were Sanger sequenced to identify correctly altered plasmids.

### Strand-specific RNA-seq quantitation

RNA-seq BAM files (GEO: GSE74660) (16) were used for gene level quantitation re-analysis. Read quantitation was performed using the featureCounts algorithm in Subread [v1.4.6-p3] software package (55) with the Ensembl v78 annotation GTF file. Settings used for quantifying the primary, reverse stranded, non-chimeric reads were as follows: -t exon -g gene_id –primary -C -s 2. Counts for genes having at least 10 counts-per-million (CPM) in all replicates of any timepoint were normalized with the TMM algorithm in the edgeR [v3.16.5] package (56) in the R [v3.3.3] environment (www.r-project.org) and exported as normalized CPM values for further analysis.

### Re-analysis of chromatin immunoprecipitation sequencing (ChIP-seq)

Raw sequencing reads for CRX (32) and NRL (33) were trimmed using trimmomatic [v0.36] (57) with parameters ‘SLIDINGWINDOW:4:15 MINLEN:24 CROP:36’. Reads were then aligned to mm10 using BWA [V0.7.15] (58) backtrack algorithm. Alignments from BWA were filtered using samtools [v1.5] (59) with following parameters’-b -q 20 -F 1796’. Duplicate alignments were then pruned using MarkDuplicates utility in Picard [v2.9.2] (http://broadinstitute.github.io/picard/). MACS2[v2.1.1] (60) with parameters‘-f BAM -g hs -B -q 0.01’ was used to identify peaks. Transcript level ChIP-seq scores were computed using a weighted sum of peak intensities metric (61). For every transcript, a region starting upstream 1,000,00 bases and downstream 10,000 bases from TSS was considered. For every peak in this region, the absolute distance (d_k_) from peak summit to TSS and pileup scores (g_k_) as computed by MACS2 were included in the formula. The d_0_ constant in metric was set to 10,000.

The distribution and density plots for ChIP-seq signals were computed and plotted using Danpos tool (62). Briefly, the dpos utility with default parameters in danpos was run on the aligned files to generate the .wig files. The profile utility in danpos was used on all promoter regions (10 kb upstream and downstream of the TSS) of candidate genes to plot the ChIP-seq signal density plot. The TSS values were extracted from Ensembl V84.

### Gene-level candidate filtering

R Studio and custom code was used to filter genes from the above large datasets. Replicates from all RNA sequencing were averaged and used for further analysis. For expression, at least
1 of 6 rod developmental time points were required to reach a minimum of 25 CPM. To identify increases in gene expression in rods, all combinations of early (P2, 4, and 6) versus late (P10, 14, and 28) were calculated; any one of these needed to reach the minimum two fold change difference. In most cases, all reached this threshold. For rod versus cone-like comparisons, each late developmental stage was compared (e.g. P10 rods versus P10 cone-like cells); a two fold change was required for any one comparison. Only genes found to have NRL and CRX ChIP-sequencing peaks (as above) were kept. We identified 719 total genes that met all of these criteria.

### Quantification and statistical analysis

R Studio and custom ggplot codes were used to generate all graphs. Prism v7 used for all statistical analysis. For all Figures and S1C, we used the Kruskal-Wallis one-way analysis of variance test with Dunn’s multiple comparison test. For Figures 2 - 5: we tested ≥25 total terminals for each construct (see Table S1); in our RNAi screen, each spherule is an independent biological replicate. Control measurements were found to have high concordance between litters and individuals, and between injected/uninjected mice, so controls were not paired with each RNAi target or rescue. When controls or RNAi knockdown data is reproduced in the original screen and rescue, these data are identical. In all cases, gene names in bold indicate significant deviation from appropriate controls with significance level as follows: *p<0.0332, **p<0.0021, ***p<0.0002, ****p<0.0001. For Figure 2B, Pearson’s correlation coefficient was used to test independence of three variables.

### Protein-protein interaction networks (PPINs)

For the presynapse associated protein interaction map, we created a network of 719 shortlisted candidates (as described above) and their 1° neighbors. *Mus musculus* protein-protein interaction data from STRING (37) was used for this analysis, where a minimum score of 700 was set to keep only high confidence interactions. This large network was mined using functions in the igraph (v1.0.1) R package, to discover network communities, and to trace all shortest paths and intermediate nodes connecting every pair of experimentally verified genes. Furthermore, shortest paths were selected to contain only those connecting via one other spherule morphology associated node. Shortest paths were also dropped if they traversed a “negative” node, representing negative genes in our RNAi screen. The presynapse associated network was created from nodes and edges that were part of filtered shortest paths that qualified the aforementioned threshold. The final network contained 71 nodes and 153 edges, derived from 96 shortest paths. All analyses were performed with custom perl and R scripts, unless mentioned otherwise. Networks were visualized using Cytoscape (v3.6) (63). GO analysis was performed using clusterProfiler (64).

## Acknowledgments

We are grateful to Robert Fariss and Wei Li for advice and Suja Hiryanna for AAV production. We thank Amal Alsufayni, Christine Park, and Kristen Mollura for technical assistance and acknowledge Andrew Smith, Freekje van Asten, and Lina Zelinger for comments on this manuscript. This research was supported by Intramural Research program of the National Eye Institute (EY000450, EY000474) and utilized the high-performance computational capabilities of the Biowulf Linux cluster at NIH (http://biowulf.nih.gov).

## Author contributions

Conceptualization: D.T.W., S.Y.K. and A.S.; Methodology: D.T.W., S.Y.K.; Investigation and Validation: D.T.W., H.F., P.H.; Software and Analysis: A.M., D.T.W., M.J.B, V.C.; Resources: W.Y., Z.W.; Writing – Original Draft: D.T.W., A.M., A.S.; Writing – Review & Editing: all authors; Supervision and Project Administration: A.S.; Funding Acquisition: A.S.

## References

1. Azevedo FA, Carvalho LR, Grinberg LT, Farfel JM, Ferretti RE, Leite RE, et al. Equal numbers of neuronal and nonneuronal cells make the human brain an isometrically scaled-up primate brain. J Comp Neurol. 2009;513(5):532–41.

2. Pakkenberg B, Pelvig D, Marner L, Bundgaard MJ, Gundersen HJ, Nyengaard JR, et al. Aging and the human neocortex. Exp Gerontol. 2003;38(1-2):95–9.

3. DeFelipe J. From the connectome to the synaptome: an epic love story. Science. 2010;330(6008):1198–201.

4. de la Torre-Ubieta L, Bonni A. Transcriptional regulation of neuronal polarity and morphogenesis in the mammalian brain. Neuron. 2011;72(1):22–40.

5. Lein E, Borm LE, Linnarsson S. The promise of spatial transcriptomics for neuroscience in the era of molecular cell typing. Science. 2017;358(6359):64–9.

6. Levine M, Davidson EH. Gene regulatory networks for development. Proc Natl Acad Sci U S A. 2005;102(14):4936–42.

7. Bonev B, Mendelson Cohen N, Szabo Q, Fritsch L, Papadopoulos GL, Lubling Y, et al. Multiscale 3D Genome Rewiring during Mouse Neural Development. Cell. 2017;171(3):557–72 e24.

8. Kittelmann S, Buffry AD, Franke FA, Almudi I, Yoth M, Sabaris G, et al. Gene regulatory network architecture in different developmental contexts influences the genetic basis of morphological evolution. PLoS Genet. 2018;14(5):e1007375.

9. Sanes JR, Zipursky SL. Design principles of insect and vertebrate visual systems. Neuron. 2010;66(1):15–36.

10. Hoon M, Okawa H, Della Santina L, Wong RO. Functional architecture of the retina: development and disease. Prog Retin Eye Res. 2014;42:44–84.

11. Masland RH. The neuronal organization of the retina. Neuron. 2012;76(2):266–80.

12. Lamb TD, Collin SP, Pugh EN, Jr. Evolution of the vertebrate eye: opsins, photoreceptors, retina and eye cup. Nat Rev Neurosci. 2007;8(12):960–76.

13. Luo DG, Xue T, Yau KW. How vision begins: an odyssey. Proc Natl Acad Sci U S A. 2008;105(29):9855–62.

14. Hunt DM, Peichl L. S cones: Evolution, retinal distribution, development, and spectral sensitivity. Vis Neurosci. 2014;31(2):115–38.

15. Lamb TD. Evolution of vertebrate retinal photoreception. Philos Trans R Soc Lond B Biol Sci. 2009;364(1531):2911–24.

16. Kim JW, Yang HJ, Brooks MJ, Zelinger L, Karakulah G, Gotoh N, et al. NRL-Regulated Transcriptome Dynamics of Developing Rod Photoreceptors. Cell Rep. 2016;17(9):2460–73.

17. Dowling JE, Boycott BB. Organization of the primate retina: electron microscopy. Proc R Soc Lond B Biol Sci. 1966;166(1002):80–111.

18. Kolb H. Organization of the outer plexiform layer of the primate retina: electron microscopy of Golgi-impregnated cells. Philos Trans R Soc Lond B Biol Sci. 1970;258(823):261–83.

19. Sterling P, Matthews G. Structure and function of ribbon synapses. Trends Neurosci. 2005;28(1):20–9.

20. Haverkamp S, Grunert U, Wassle H. The cone pedicle, a complex synapse in the retina. Neuron. 2000;27(1):85–95.

21. Rao-Mirotznik R, Harkins AB, Buchsbaum G, Sterling P. Mammalian rod terminal: architecture of a binary synapse. Neuron. 1995;14(3):561–9.

22. Mears AJ, Kondo M, Swain PK, Takada Y, Bush RA, Saunders TL, et al. Nrl is required for rod photoreceptor development. Nat Genet. 2001;29(4):447–52.

23. Oh EC, Khan N, Novelli E, Khanna H, Strettoi E, Swaroop A. Transformation of cone precursors to functional rod photoreceptors by bZIP transcription factor NRL. Proc Natl Acad Sci U S A. 2007;104(5):1679–84.

24. Brzezinski JA, Reh TA. Photoreceptor cell fate specification in vertebrates. Development. 2015;142(19):3263–73.

25. Swaroop A, Kim D, Forrest D. Transcriptional regulation of photoreceptor development and homeostasis in the mammalian retina. Nat Rev Neurosci. 2010;11(8):563–76.

26. Blanks JC, Adinolfi AM, Lolley RN. Synaptogenesis in the photoreceptor terminal of the mouse retina. J Comp Neurol. 1974;156(1):81–93.

27. Strettoi E, Mears AJ, Swaroop A. Recruitment of the rod pathway by cones in the absence of rods. J Neurosci. 2004;24(34):7576–82.

28. Daniele LL, Lillo C, Lyubarsky AL, Nikonov SS, Philp N, Mears AJ, et al. Cone-like morphological, molecular, and electrophysiological features of the photoreceptors of the Nrl knockout mouse. Invest Ophthalmol Vis Sci. 2005;46(6):2156–67.

29. Sarin S, Zuniga-Sanchez E, Kurmangaliyev YZ, Cousins H, Patel M, Hernandez J, et al. Role for Wnt Signaling in Retinal Neuropil Development: Analysis via RNA-Seq and In Vivo Somatic CRISPR Mutagenesis. Neuron. 2018;98(1):109–26 e8.

30. Carter-Dawson LD, LaVail MM. Rods and cones in the mouse retina. I. Structural analysis using light and electron microscopy. J Comp Neurol. 1979;188(2):245–62.

31. Li S, Mitchell J, Briggs DJ, Young Jk, Long SS, Fuerst PG. Morphological Diversity of the Rod Spherule: A Study of Serially Reconstructed Electron Micrographs. PLoS One. 2016;11(3):e0150024.

32. Corbo JC, Lawrence KA, Karlstetter M, Myers CA, Abdelaziz M, Dirkes W, et al. CRX ChIP-seq reveals the cis-regulatory architecture of mouse photoreceptors. Genome Res. 2010;20(11):1512–25.

33. Hao H, Kim DS, Klocke B, Johnson KR, Cui K, Gotoh N, et al. Transcriptional regulation of rod photoreceptor homeostasis revealed by in vivo NRL targetome analysis. PLoS Genet. 2012;8(4):e1002649.

34. Sanuki R, Watanabe S, Sugita Y, Irie S, Kozuka T, Shimada M, et al. Protein-4.1G-Mediated Membrane Trafficking Is Essential for Correct Rod Synaptic Location in the Retina and for Normal Visual Function. Cell Rep. 2015;10:796–808.

35. Adams ME, Kramarcy N, Krall SP, Rossi SG, Rotundo RL, Sealock R, et al. Absence of alpha-syntrophin leads to structurally aberrant neuromuscular synapses deficient in utrophin. J Cell Biol. 2000; 150(6):1385–98.

36. Yu W, Mookherjee S, Chaitankar V, Hiriyanna S, Kim JW, Brooks M, et al. Nrl knockdown by AAV-delivered CRISPR/Cas9 prevents retinal degeneration in mice. Nat Commun. 2017;8:14716.

37. Szklarczyk D, Morris JH, Cook H, Kuhn M, Wyder S, Simonovic M, et al. The STRING database in 2017: quality-controlled protein-protein association networks, made broadly accessible. Nucleic Acids Res. 2017;45(D1):D362–D8.

38. Swanson LW, Bota M. Foundational model of structural connectivity in the nervous system with a schema for wiring diagrams, connectome, and basic plan architecture. Proc Natl Acad Sci U S A. 2010;107(48):20610–7.

39. Regus-Leidig H, Tom Dieck S, Specht D, Meyer L, Brandstatter JH. Early steps in the assembly of photoreceptor ribbon synapses in the mouse retina: the involvement of precursor spheres. J Comp Neurol. 2009;512(6):814–24.

40. Lagnado L, Schmitz F. Ribbon Synapses and Visual Processing in the Retina. Annu Rev Vis Sci. 2015;1:235–62.

41. Maxeiner S, Luo F, Tan A, Schmitz F, Sudhof TC. How to make a synaptic ribbon: RIBEYE deletion abolishes ribbons in retinal synapses and disrupts neurotransmitter release. EMBO J. 2016;35(10):1098–114.

42. Regus-Leidig H, Fuchs M, Lohner M, Leist SR, Leal-Ortiz S, Chiodo VA, et al. In vivo knockdown of Piccolino disrupts presynaptic ribbon morphology in mouse photoreceptor synapses. Front Cell Neurosci. 2014;8:259.

43. Cao Y, Sarria I, Fehlhaber KE, Kamasawa N, Orlandi C, James KN, et al. Mechanism for Selective Synaptic Wiring of Rod Photoreceptors into the Retinal Circuitry and Its Role in Vision. Neuron. 2015;87(6):1248–60.

44. Sato S, Omori Y, Katoh K, Kondo M, Kanagawa M, Miyata K, et al. Pikachurin, a dystroglycan ligand, is essential for photoreceptor ribbon synapse formation. Nat Neurosci. 2008;11(8):923–31.

45. Wang Y, Fehlhaber KE, Sarria I, Cao Y, Ingram NT, Guerrero-Given D, et al. The Auxiliary Calcium Channel Subunit alpha2delta4 Is Required for Axonal Elaboration, Synaptic Transmission, and Wiring of Rod Photoreceptors. Neuron. 2017;93(6):1359–74 e6.

46. Matsuda T, Cepko CL. Electroporation and RNA interference in the rodent retina in vivo and in vitro. Proc Natl Acad Sci U S A. 2004;101(1):16–22.

47. de Melo J, Blackshaw S. In Vivo Electroporation of Developing Mouse Retina. Methods Mol Biol. 2018;1715:101–11.

48. Mitton KP, Swain PK, Chen S, Xu S, Zack DJ, Swaroop A. The leucine zipper of NRL interacts with the CRX homeodomain. A possible mechanism of transcriptional synergy in rhodopsin regulation. J Biol Chem. 2000;275(38):29794–9.

49. Assawachananont J, Kim SY, Kaya KD, Fariss R, Roger JE, Swaroop A. Cone-rod homeobox CRX controls presynaptic active zone formation in photoreceptors of mammalian retina. Hum Mol Genet. 2018;27(20):3555–67.

50. Yoshimatsu T, D’Orazi FD, Gamlin CR, Suzuki SC, Suli A, Kimelman D, et al. Presynaptic partner selection during retinal circuit reassembly varies with timing of neuronal regeneration in vivo. Nat Commun. 2016;7:10590.

51. Akimoto M, Cheng H, Zhu D, Brzezinski JA, Khanna R, Filippova E, et al. Targeting of GFP to newborn rods by Nrl promoter and temporal expression profiling of flow-sorted photoreceptors. Proc Natl Acad Sci U S A. 2006;103(10):3890–5.

52. Khani SC, Pawlyk BS, Bulgakov OV, Kasperek E, Young JE, Adamian M, et al. AAV-mediated expression targeting of rod and cone photoreceptors with a human rhodopsin kinase promoter. Invest Ophthalmol Vis Sci. 2007;48(9):3954–61.

53. Grimm D, Zhou S, Nakai H, Thomas CE, Storm TA, Fuess S, et al. Preclinical in vivo evaluation of pseudotyped adeno-associated virus vectors for liver gene therapy. Blood. 2003;102(7):2412–9.

54. Schneider CA, Rasband WS, Eliceiri KW. NIH Image to ImageJ: 25 years of image analysis. Nat Methods. 2012;9(7):671–5.

55. Liao Y, Smyth GK, Shi W. featureCounts: an efficient general purpose program for assigning sequence reads to genomic features. Bioinformatics. 2014;30(7):923–30.

56. Robinson MD, McCarthy DJ, Smyth GK. edgeR: a Bioconductor package for differential expression analysis of digital gene expression data. Bioinformatics. 2010;26(1):139–40.

57. Bolger AM, Lohse M, Usadel B. Trimmomatic: a flexible trimmer for Illumina sequence data. Bioinformatics. 2014;30(15):2114–20.

58. Li H, Durbin R. Fast and accurate short read alignment with Burrows-Wheeler transform. Bioinformatics. 2009;25(14):1754–60.

59. Li H, Handsaker B, Wysoker A, Fennell T, Ruan J, Homer N, et al. The Sequence Alignment/Map format and SAMtools. Bioinformatics. 2009;25(16):2078–9.

60. Zhang Y, Liu T, Meyer CA, Eeckhoute J, Johnson DS, Bernstein BE, et al. Model-based analysis of ChIP-Seq (MACS). Genome Biol. 2008;9(9):R137.

61. Ouyang Z, Zhou Q, Wong WH. ChIP-Seq of transcription factors predicts absolute and differential gene expression in embryonic stem cells. Proc Natl Acad Sci U S A. 2009;106(51):21521–6.

62. Chen K, Xi Y, Pan X, Li Z, Kaestner K, Tyler J, et al. DANPOS: dynamic analysis of nucleosome position and occupancy by sequencing. Genome Res. 2013;23(2):341–51.

63. Shannon P, Markiel A, Ozier O, Baliga NS, Wang JT, Ramage D, et al. Cytoscape: a software environment for integrated models of biomolecular interaction networks. Genome Res. 2003;13(11):2498–504.

64. Yu G, Wang LG, Han Y, He QY. clusterProfiler: an R package for comparing biological themes among gene clusters. OMICS. 2012;16(5):284–7.

